# An anti-aggregation region of the SGS3 N-terminal IDR is essential for secondary siRNA biogenesis

**DOI:** 10.64898/2026.01.02.697351

**Authors:** Yuji Fujimoto, Yuriki Sakurai, Ryosuke Kowada, Keisuke Shoji, Manabu Yoshikawa, Hiro-oki Iwakawa

## Abstract

Secondary siRNA biogenesis amplifies small RNA signals from target transcripts and plays a pivotal role in plant development and defense responses. The RNA-binding protein SGS3 is essential for this pathway, recruiting RNA-dependent RNA polymerase 6 (RDR6) to Argonaute–small RNA–bound targets. The N-terminal intrinsically disordered region (IDR) of SGS3, which contains a prion-like domain (PrLD), has been reported to drive liquid–liquid phase separation, forming siRNA bodies, and to be required for secondary siRNA production. However, the molecular mechanism by which the N-terminal IDR contributes to secondary siRNA production remains unclear. Here, using in vitro reconstitution and in planta assays, we show that the N-terminal IDR comprises two functional modules: the PrLD and a negatively charged region (NCR). The PrLD is required for phase separation and siRNA body formation but is dispensable for secondary siRNA production. In contrast, mutations in the NCR caused SGS3 to form abnormally large cytoplasmic assemblies and markedly impaired secondary siRNA production. These results suggest that the N-terminal IDR helps maintain SGS3 in a functional, soluble state that supports efficient secondary siRNA biogenesis.

## Introduction

Small RNAs, including microRNAs (miRNAs) and small interfering RNAs (siRNAs), induce RNA silencing, a sequence-specific gene regulation mechanism involved in numerous biological processes (Ghildiyal & Zamore, 2009). To silence target genes, small RNAs must assemble into an effector complex known as RNA-induced silencing complex (RISC), which contains an Argonaute (AGO) protein (Iwakawa & Tomari, 2022). RISC recognizes target RNAs based on sequence complementarity and suppresses their expression through RNA cleavage or translational repression (Iwakawa & Tomari, 2022). In some organisms, including plants, fungi, and worms, specific RISCs recruit RNA-dependent RNA polymerases (RDRs) to their target RNAs (Baulcombe, 2007; Dang *et al*, 2011). RDRs synthesize double-stranded RNAs (dsRNAs) from these targets, which are subsequently processed into secondary siRNAs by Dicer or Dicer-like (DCL) proteins (Sijen *et al*, 2001). Such secondary siRNAs are key regulators of development, stress adaptation, reproductive processes, antiviral immunity, genome stability, and transgene expression efficiency (Bologna & Voinnet, 2014; Liu *et al*, 2020).

Plants possess two major pathways for secondary siRNA biogenesis: one that suppresses foreign or self-replicating nucleic acids such as viruses and transposons, and another that regulates endogenous gene expression (Baulcombe, 2007; Liu *et al*, 2020; Sanan-Mishra *et al*, 2021; Baulcombe, 2022). The former, known as sense-transgene–induced post-transcriptional gene silencing (S-PTGS), operates without any endogenous primary small RNA trigger. In contrast, the latter pathway is initiated when specific miRNAs trigger the production of phased secondary siRNAs (phasiRNAs) from target transcripts encoded at PHAS loci. Among these, trans-acting siRNAs (tasiRNAs) act on genes distinct from their loci of origin and are produced from TAS loci (Liu *et al*, 2020). In *Arabidopsis thaliana*, eight TAS loci have been identified: *TAS1a/b/c*, *TAS2*, *TAS3a/b/c*, and *TAS4* (Allen *et al*, 2005; Yoshikawa *et al*, 2005; Rajagopalan *et al*, 2006; Howell *et al*, 2007). Transcripts derived from *TAS1* and *TAS2* genes are recognized by the miR173–AGO1 RISC, *TAS3* transcripts by the miR390–AGO7 RISC, and *TAS4* transcripts by the miR828–AGO1 RISC. These transcripts are recognized by specific RISCs, and are converted into double-stranded RNA through RDR6 recruitment, leading to the production of tasiRNAs. RISCs that have been reported to possess the ability to recruit RDR6 are limited to AGO1–RISC loaded with a 22-nt small RNA and AGO7–RISC loaded with miR390, which specifically recognizes *TAS3* transcripts (Axtell *et al*, 2006; Chen *et al*, 2010; Cuperus *et al*, 2010). SUPPRESSOR OF GENE SILENCING 3 (SGS3) specifically associates with these RISCs and their target RNAs and, in cooperation with SILENCING DEFECTIVE 5 (SDE5), recruits RDR6 to the target mRNAs (Iwakawa *et al*, 2021; Sakurai *et al*, 2021; Yoshikawa *et al*, 2021, 2013; Fujimoto & Iwakawa, 2023). RDR6 then synthesizes dsRNA, which is processed by DCL2/3/4 into secondary siRNAs (Howell *et al*, 2007; Rajeswaran *et al*, 2012). These sequential reactions occur in membrane-less organelles termed siRNA bodies (Kumakura *et al*, 2009; Jouannet *et al*, 2012; Tan *et al*, 2023).

Many membrane-less organelles are formed through liquid–liquid phase separation (LLPS) (Musacchio, 2022), which underlies the assembly of diverse compartments that modulate biochemical reactions such as transcription, stress responses, and cell signaling (Hirose *et al*, 2023; O’Flynn & Mittag, 2021). RNA silencing pathways have also been shown to utilize LLPS to regulate reaction dynamics. For instance, in human cells, LLPS formed by human AGO2 and the scaffold protein TNRC6 (GW182) recruits the CCR4–NOT deadenylation factors, thereby accelerating mRNA deadenylation (Sheu-Gruttadauria & MacRae, 2018). In *A. thaliana*, the SERRATE (SE) protein forms dicing bodies that promote miRNA processing (Xie *et al*, 2021; Zhong *et al*, 2024). It has been reported that siRNA bodies are assembled around SGS3, which functions as a scaffold. Because factors involved in secondary siRNA biogenesis̶such as RDR6, AGO7, and SDE5—localize to siRNA bodies, these structures are thought to represent sites of secondary siRNA biogenesis (Tan *et al*, 2023; Jouannet *et al*, 2012; Kumakura *et al*, 2009).

SGS3 is a plant-specific double-stranded RNA-binding protein. SGS3 contains six known domains—an N-terminal prion-like domain (PrLD_N) within the intrinsically disordered region (IDR), a zinc-finger (ZF), an XS domain, coiled-coil 1 (CC1), PrLD_C, and coiled-coil 2 (CC2) (Figure EV. 1)—and is thought to function as a homodimer mediated by its coiled-coil domains (Elmayan *et al*, 2009; Fukunaga & Doudna, 2009). Biochemical analyses have suggested that SGS3 preferentially binds to double-stranded RNAs with a 5′ overhang through its XS domain (Fukunaga & Doudna, 2009). The PrLD is a low-complexity domain known to drive phase separation (Kato *et al*, 2012; Banani *et al*, 2017). Deletion of the N-terminal IDR containing the PrLD_N abolishes siRNA body formation and causes *Arabidopsis* to exhibit an *sgs3* mutant–like phenotype, leading to the proposal that PrLD-mediated siRNA body assembly is required for secondary siRNA production (Tan *et al*, 2023; Kim *et al*, 2021). However, it has not been directly demonstrated whether siRNA body formation itself is essential for secondary siRNA biogenesis, nor has it been examined whether functional elements within the N-terminal IDR other than the PrLD_N contribute to siRNA body formation and secondary siRNA biogenesis.

To dissect the role of the N-terminal IDR of SGS3 in secondary siRNA biogenesis, we performed both *in vitro* reconstitution and in planta functional analyses. We demonstrate that, although the PrLD_N is required for LLPS and siRNA body formation, it is dispensable for endogenous secondary siRNA accumulation. Instead, a negatively charged region (NCR) within the non-PrLD portion of the N-terminal IDR is required to prevent abnormal aggregation of SGS3 and is indispensable for endogenous secondary siRNA accumulation. This NCR may facilitate the secondary siRNA-producing activity of SGS3 by exerting an anti-aggregation function.

## Results

### The N-terminal prion-like domain of SGS3 is required for condensate formation but not for tasiRNA production *in vitro*

First, we examined whether condensate formation by SGS3 is dependent on the PrLD. Although previous studies have concluded that the PrLD located at the N-terminus of SGS3 induces its phase separation (Tan *et al*, 2023; Kim *et al*, 2021), these analyses were performed using mutants with large deletions of at least ∼200 amino acids from the N-terminal region, making it unclear whether the PrLD itself or other sequences adjacent to it are responsible for driving phase separation. To clarify this, we examined condensate formation *in vitro* using recombinant SGS3 fused to EGFP. mRNAs encoding EGFP-tagged wild-type SGS3 or its PrLD-deleted form (ΔPrLD_N–EGFP) were translated in the tobacco cell-free system (Komoda *et al*, 2004) for 6 h, followed by fluorescence microscopy (Figure 1A). Expression of wild-type SGS3–EGFP led to the formation of numerous condensates, whereas EGFP alone or ΔPrLD_N–EGFP—despite comparable expression levels—did not form condensates (Figure 1B, C, D). Collectively, these *in vitro* results demonstrate that the N-terminal PrLD of SGS3 is required for condensate formation but is dispensable for tasiRNA biogenesis.

**Figure 1.**
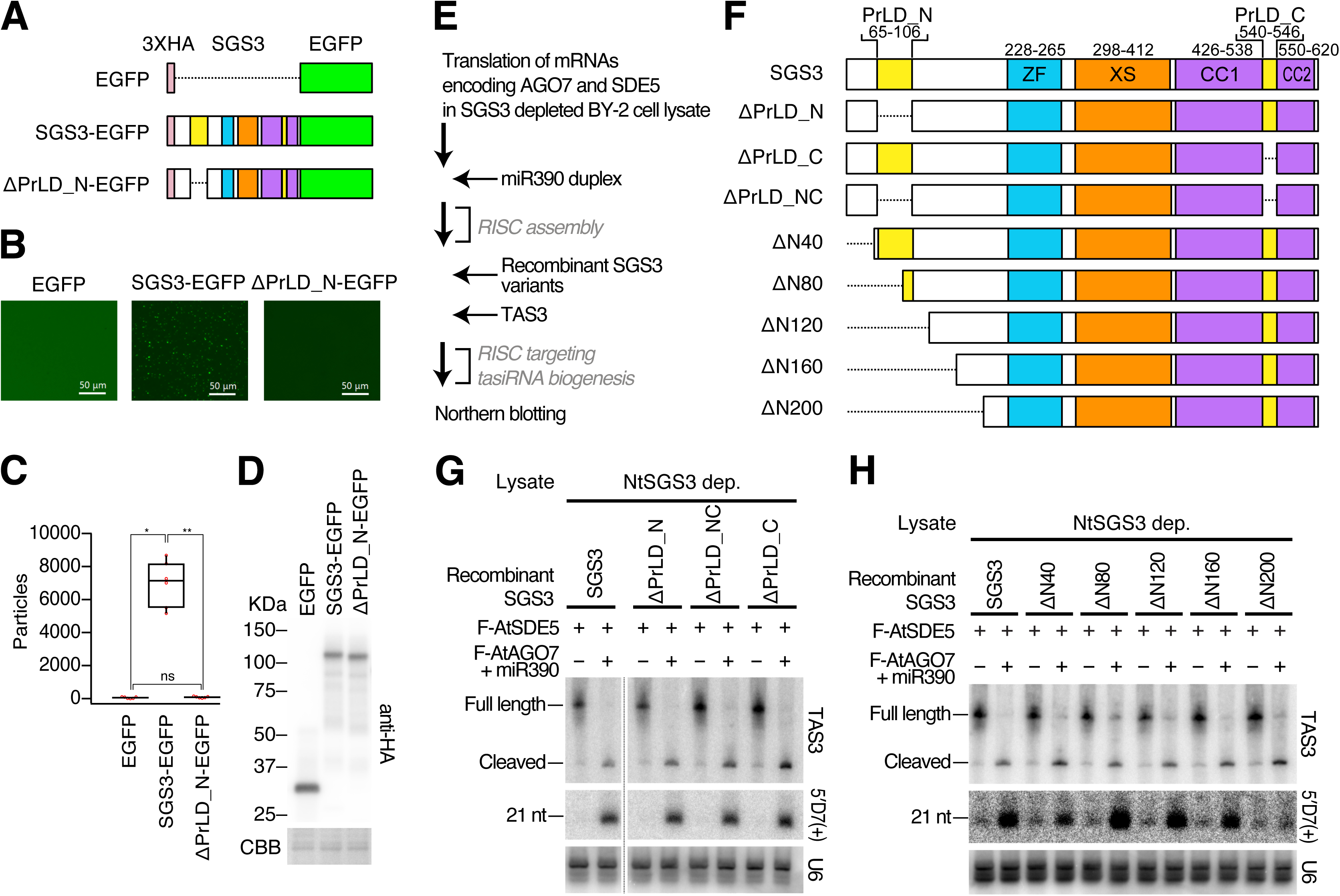
The PrLD contained within the N-terminal IDR of SGS3 is not essential for secondary siRNA production. (A) Schematic representation of SGS3 constructs used for *in vitro* condensate formation. The N-terminal PrLD-deletion mutant and wild-type (WT) SGS3 were fused to EGFP at their C termini. (B) Fluorescence microscopy images of condensates formed by EGFP-tagged SGS3 proteins shown in (A). (C) Quantification of the number of condensates observed in (B). The mean values ± SD from four independent experiments are shown. Bonferroni-corrected p values from two-sided paired t tests are as follows: *p = 0.00000087 and **p = 0.00000083. (D) Western blot analysis of protein levels in the samples used for the observations in (B). (E) Schematic workflow of *in vitro* reconstitution of TAS3 tasiRNA biogenesis assay that we recently developed. First, we added mRNAs encoding AGO7 and SDE5 in SGS3 depleted BY-2 lysate. Next, miR390 duplex was added to form AGO7-RISC. Recombinant SGS3 variants and *TAS3* transcripts were then added to trigger tasiRNA biogenesis. (F) Schematic representation of the SGS3 domain organization and mutant design. SGS3 consists of 625 amino acids, and the numbers indicate the positions from the N-terminus. Dashed lines represent the deleted regions. (G) *In vitro* tasiRNA biogenesis with SGS3 mutants lacking the one or both PrLDs. *TAS3* transcripts and 5′ D7 (+) tasiRNA were detected by northern blotting. U6 spliceosomal RNA was used as a loading control. (H)*In vitro* tasiRNA biogenesis with SGS3 N-terminal deletion mutants. *TAS3* transcripts and 5′ D7 (+) tasiRNA were detected by northern blotting. U6 spliceosomal RNA was used as a loading control.

Next, we examined whether the PrLD is required for secondary siRNA production. For this purpose, we used a recently developed *in vitro* reconstitution system that recapitulates secondary siRNA biogenesis using a lysate derived from *Nicotiana tabacum* BY-2 suspension cells (Sakurai *et al*, 2021; Iki *et al*, 2010; Komoda *et al*, 2004).

This cell-free system can produce tasiRNAs *in vitro* upon the addition of mRNAs encoding AGO7 and SDE5, along with the miR390 duplex and its target *TAS3* transcript. Although the lysate contains endogenous SGS3, RDR6, and Dicer-like (DCL) proteins at levels sufficient to support tasiRNA production, their individual functions can be dissected by antibody-mediated selective depletion followed by complementation. To analyze domain-specific requirements of SGS3, endogenous *Nicotiana tabacum* SGS3 (NtSGS3) was removed from the lysate by immunodepletion, and recombinant *Arabidopsis thaliana* SGS3 (AtSGS3) or its mutants were added back (Figure 1E). The recombinant wild-type AtSGS3 successfully restored TAS3 tasiRNA production. In contrast, deletion mutants lacking the ZF, XS, or CC domains (ΔZF, ΔXS, ΔCC) (Kumakura *et al*, 2009), as well as the E500K point mutant (Adenot *et al*, 2006) failed to support tasiRNA biogenesis, consistent with in planta observations (Figure EV. 2A, B). Notably, these inactive mutants also exhibited dominant-negative effects in lysates containing endogenous NtSGS3 (mock-depleted lysates; Figure EV. 2A). Together, these results confirm that the *in vitro* reconstitution system faithfully reproduces SGS3-dependent tasiRNA biogenesis observed in vivo.

To further investigate the relationship between siRNA body formation and tasiRNA production, we next focused on the PrLD of SGS3, which constitutes the core of siRNA bodies, and examined whether the PrLD itself is involved in tasiRNA biogenesis. To this end, we generated three SGS3 mutants—ΔPrLD_N (Δ65–106), ΔPrLD_C (Δ540–546), and ΔPrLD_NC (lacking both regions)—and tested their ability to restore tasiRNA production in the cell-free assay (Figure 1F). All three mutants supported tasiRNA synthesis at levels comparable to wild-type SGS3 (Figure 1G), indicating that the PrLD is not required for TAS3 tasiRNA biogenesis *in vitro*.

In a previous report, *A. thaliana* expressing an SGS3 variant lacking approximately 200 amino acids at the N-terminus exhibited a downward-curled leaf phenotype similar to that of *sgs3* mutants (Peragine *et al*, 2004). Although secondary siRNA production was not directly examined in that study, if this phenotype arises from a defect in secondary siRNA biogenesis, it would indicate that deletion of the 200–amino-acid region leads to impaired secondary siRNA production. Given our finding that the PrLD is not required for endogenous secondary siRNA biogenesis, these observations suggest that the N-terminal 200 amino acids of SGS3 contain additional functional element(s), distinct from the PrLD, that are essential for secondary siRNA production. To test this possibility, we generated five recombinant AtSGS3 truncation mutants—ΔN40, ΔN80, ΔN120, ΔN160, and ΔN200—harboring sequential deletions of 40–200 amino acids from the N terminus, and evaluated their ability to support tasiRNA synthesis in the cell-free assay (Figure 1B). Deletions up to 160 amino acids did not affect tasiRNA production (ΔN160), whereas deletion of 200 amino acids markedly reduced tasiRNA synthesis (ΔN200) (Figure 1H). These findings indicate that, while the PrLD indeed facilitates siRNA body formation driven by LLPS, the N-terminal 160 amino acids containing the PrLD are not required for secondary siRNA production itself. Instead, the downstream region between residues 160 and 200 appears to play an indispensable role in this process.

### The N-terminal prion-like domain of SGS3 is required for siRNA body formation but not for phasiRNA/tasiRNA biogenesis *in vivo*

Our *in vitro* reconstitution experiments demonstrated that the N-terminal PrLD of SGS3 is required for condensate formation but dispensable for tasiRNA production. However, *in vitro* reactions occur under conditions of reduced macromolecular crowding and lack the diverse intracellular compartments present in living cells. In the cytoplasm, SGS3 likely encounters various phase-separated granules, such as processing bodies (Iwasaki *et al*, 2007; Xu *et al*, 2007) and stress granules (Jouannet *et al*, 2012), which may influence the assembly or function of siRNA bodies. Therefore, it remained possible that the PrLD contributes to SGS3 function in the more complex *in vivo* environment, either by mediating interactions with other condensates or by fine-tuning its phase behavior. To test this hypothesis, we next analyzed the functional relevance of the N-terminal region of SGS3 in *A. thaliana*.

To evaluate the *in vivo* function of the SGS3 N-terminal region, we generated transgenic *sgs3-11* plants expressing FLAG-tagged SGS3, its N-terminal mutants, or Firefly luciferase (negative control) under the constitutive 35S promoter (35S:SGS3, 35S:ΔN160, 35S:ΔN200, 35S:ΔPrLD_N, and 35S:FLUC) (Figure 2A). Because the functional activity of SGS3 may depend on its expression level, we generated both high-expression (“_H”) and low-expression (“_L”) T1 transgenic progeny for each construct and used these plants for subsequent analyses (Figure 2B). Immunoblot analysis using an antibody against the N-terminus of AtSGS3 revealed that the low-expression progeny accumulated less than 5% of endogenous AtSGS3 in Col-0, whereas the high-expression progeny accumulated approximately 20-fold higher levels (Figure 2C).

**Figure 2.**
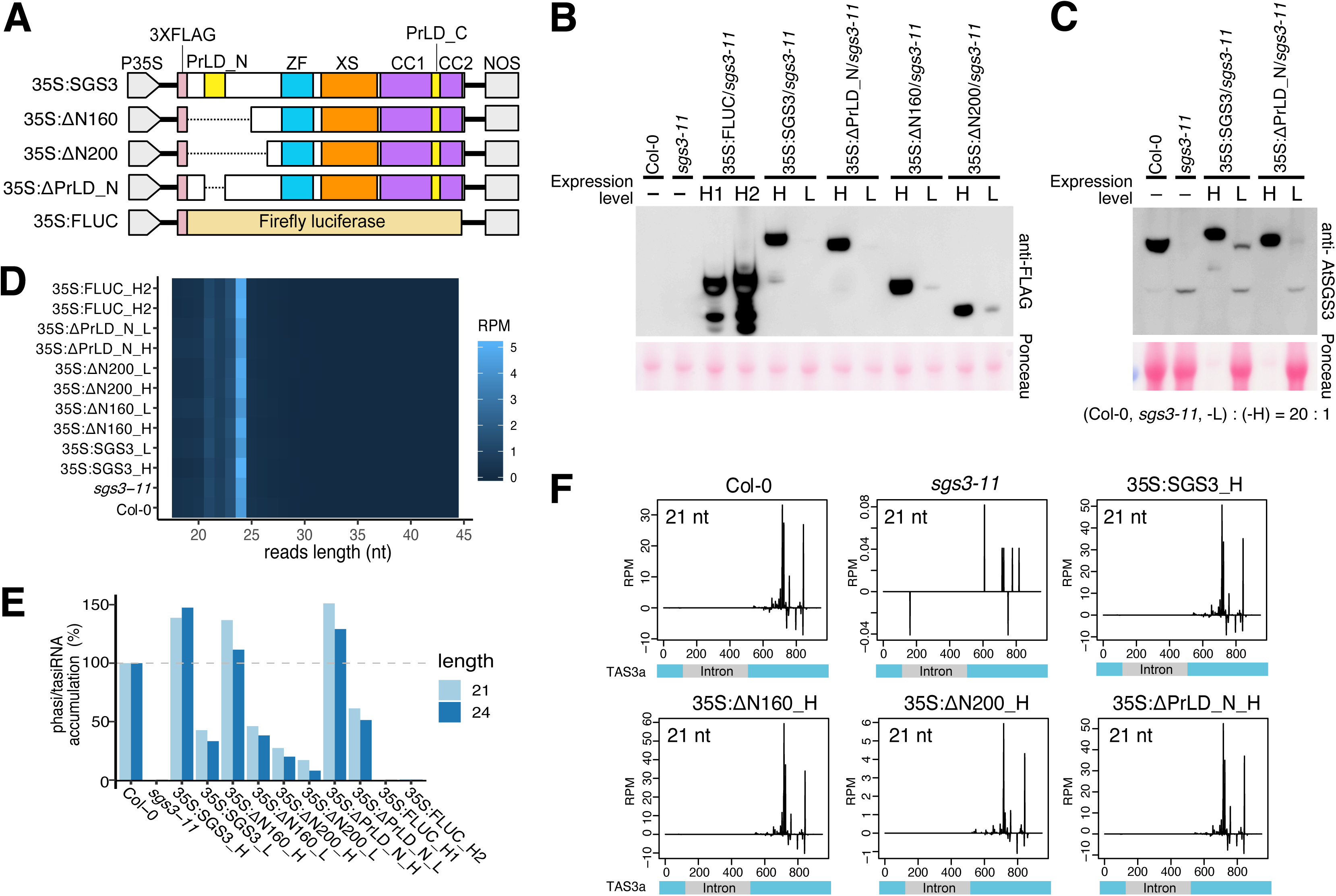
Contribution of the N-terminal region of SGS3 to tasiRNA production in *Arabidopsis* plants. (A) Schematic representation of the constructs used for *A. thaliana* transformation. Gene expression was driven by the CaMV 35S promoter and terminated by the NOS terminator. Firefly luciferase (FLUC) was used as a negative control. (B) Western blot analysis of FLAG-tagged proteins in *Arabidopsis sgs3-11* T1 progeny transformed with WT SGS3, SGS3 mutant, or FLUC control constructs. Detection was performed using an anti-FLAG antibody, and Ponceau staining was used as a loading control. In the lane labels, “H” indicates a progeny with high protein accumulation (High), and “L” indicates a progeny with low accumulation (Low). (C) Western blot analysis of SGS3 protein accumulation in *Arabidopsis* Col-0 plant and *sgs3-11* T1 progeny. Detection was performed using an anti-AtSGS3 antibody, and Ponceau staining was used as a loading control. To allow comparison between endogenous SGS3 (Col-0) and the introduced SGS3 proteins, samples from Col-0 and low-expression (“L”) progeny were loaded with the same amount of total protein, whereas high-expression (“H”) progeny were loaded with one-twentieth of that amount, as reflected in the Ponceau-stained bands. (D–F) Results of small RNA sequencing using total RNA extracted from *A. thaliana*. (D) Heat map of small RNA length distribution, with lighter cyan indicating higher abundance. (E) Relative accumulation of 21-nt and 24-nt PHAS/TAS-derived small RNAs in Col-0 and each complemented *sgs3-11* progeny. Values are shown as bar graphs, with the abundance in Col-0 set to 100 (RPM) for each size class. (F) Profiles of 21-nt TAS3-derived small RNAs mapped along the *TAS3a* genomic region, plotted by the 5′ ends of small RNA reads (RPM).

We then analyzed the small RNA populations in these transgenic progeny by performing small RNA sequencing of flower buds and comparing the accumulation of siRNAs derived from *PHAS/TAS* loci. First, the overall length distribution of small RNAs (18–44 nt) was similar among all genotypes, including *sgs3-11*, with peaks at 24 nt followed by 21 nt and 23 nt (Figure 2D). Thus, deletion or mutation of SGS3 did not affect the general length distribution of small RNAs.

As expected, *sgs3-11* mutants exhibited a near-complete loss of phasi/tasiRNA accumulation (Figure 2E), consistent with previous reports that SGS3 is essential for endogenous secondary siRNA production (Vazquez *et al*, 2004; Allen *et al*, 2005; Yoshikawa *et al*, 2005). In contrast, transgenic plants expressing wild-type SGS3, ΔN160, or ΔPrLD_N restored phasi/tasiRNA accumulation to levels comparable to or slightly higher than Col-0. In the high-expression progeny (“_H”), these constructs yielded approximately 140% of Col-0 tasiRNA levels, whereas in the low-expression progeny (“_L”), 35S:SGS3_L and 35S:ΔN160_L restored tasiRNA accumulation to ∼50% of Col-0 levels, and 35S:ΔPrLD_N_L to ∼60% (Figure 2E). By contrast, 35S:ΔN200_H and 35S:ΔN200_L accumulated only ∼27% and ∼17% of Col-0 tasiRNA levels, respectively.

Notably, although SGS3 protein abundance in the high-expression progeny was approximately 20-fold greater than that of endogenous SGS3 in Col-0, tasiRNA accumulation increased only about 1.4-fold (Figure 2B, C, E). Conversely, when SGS3 abundance was reduced to roughly one-twentieth of wild-type levels in the low-expression progeny, tasiRNA levels still remained at ∼50% of Col-0. These data indicate that tasiRNA biogenesis is relatively insensitive to SGS3 dosage and can be maintained across a broad range of SGS3 expression levels. Thus, tasiRNA production is robust to fluctuations in SGS3 abundance, and deletions up to residue 160, including the PrLD, do not impair phasi/tasiRNA production, whereas the region between residues 160 and 200 is critical for efficient secondary siRNA biogenesis.

We further examined whether deletions in the N-terminal IDR affected the production or phasing patterns of tasiRNAs triggered by AGO1 (*TAS1* and *TAS2*) and AGO7 (*TAS3*). As expected, tasiRNA production was almost completely lost in *sgs3-11*, while transgenic expression of wild-type SGS3 or the N-terminal deletion mutants restored both normal phasing and abundance (Figure 2F). Consistent with total tasiRNA levels, 35S:SGS3, 35S:ΔPrLD_N, and 35S:ΔN160 restored tasiRNA accumulation to wild-type levels or higher, whereas 35S:ΔN200 remained defective.

Collectively, these *in vivo* analyses indicate that most of the N-terminal IDR of SGS3, including the PrLD, is dispensable for endogenous secondary siRNA biogenesis, whereas the proximal region adjacent to the ZF domain (residues 160–200) plays a crucial role in promoting secondary siRNA production.

### A negatively charged region within the N-terminal IDR prevents aberrant aggregation of SGS3 in plants and is essential for tasiRNA production

Our *in vitro* and *in vivo* analyses showed that deletions extending up to residue 160 of SGS3, including the PrLD, had little or no impact on phasi/tasiRNA biogenesis, whereas further deletion to residue 200 abolished tasiRNA production. Because the ΔN200 mutant failed to support tasiRNA synthesis, we hypothesized that the region between amino acid residues 160 and 200 contributes to the proper assembly or structural organization of SGS3. To investigate whether this functional defect is associated with altered subcellular behavior, we next examined the localization and morphological features of SGS3 and its N-terminal mutants, including their condensate-forming properties, in living cells.

Transient expression assays were performed in *Nicotiana benthamiana* leaves to visualize the subcellular localization of wild-type SGS3 and its mutants fused to EGFP (Figure 3A). Granule morphology was further analyzed by quantifying the roundness of SGS3-containing structures using fluorescence images. Free EGFP showed a diffuse cytoplasmic distribution, whereas wild-type SGS3 (SGS3–EGFP) formed numerous small, spherical cytoplasmic bodies, consistent with previous reports (Tan *et al*, 2023; Kumakura *et al*, 2009). ΔPrLD_N–EGFP and ΔN160–EGFP, which retained tasiRNA-producing ability, exhibited diffuse cytoplasmic fluorescence similar to EGFP and rarely formed visible condensates (Figure 3B). These observations support the conclusion that siRNA body formation per se is not essential for tasiRNA production. In contrast, the ΔN200–EGFP mutant, which was defective in tasiRNA biogenesis, formed large and irregular aggregates that were clearly distinct from normal siRNA bodies (Figure 3B). The neighboring ZF domain deletion mutant, ΔZF–EGFP, formed small, round bodies indistinguishable from SGS3–EGFP, and, consistent with a previous report (Kumakura *et al*, 2009), did not produce abnormal aggregates. These findings suggest that the region adjacent to the ZF domain within the N-terminal IDR plays an important role in suppressing aberrant aggregation of SGS3. Consistent with this idea, a prominent negatively charged region (NCR) enriched in Asp and Glu residues was identified between residues 160 and 190; this region is highly conserved among plant species (Figure EV4A).

**Figure 3.**
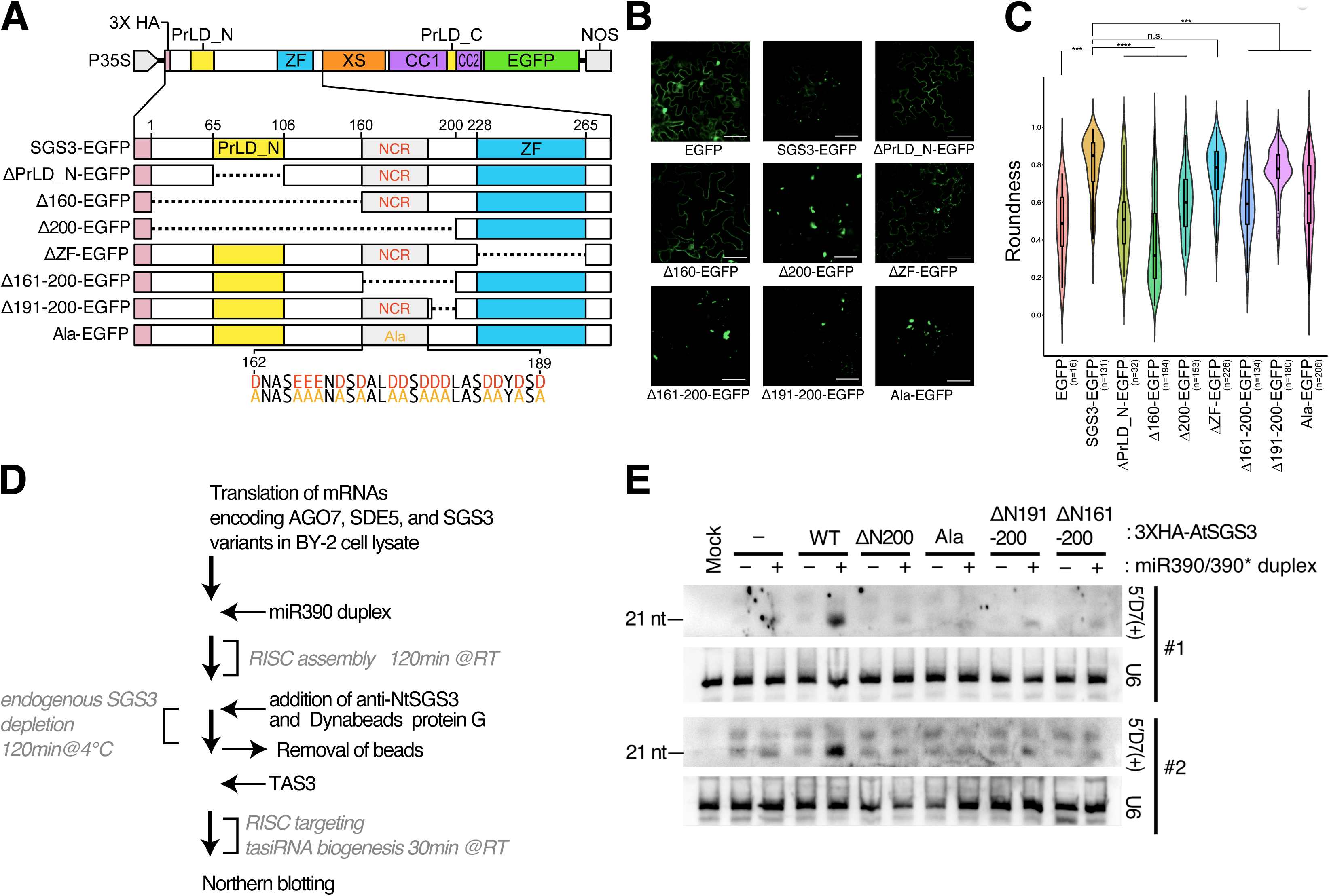
The conserved negatively charged region within the N-terminal IDR of SGS3 is essential for preventing aberrant aggregation, as well as for tasiRNA production. (A) Schematic representation of EGFP-tagged SGS3 mutant constructs used for fluorescence microscopy. Dashed lines indicate deleted regions, and numbers correspond to amino acid positions from the initiating methionine of the SGS3 open reading frame. (B) Fluorescence microscopy images showing the subcellular localization of SGS3 mutants transiently expressed in *N. benthamiana*. The white scale bar in each image represents 50 µm. (C) Quantitative analysis of the roundness of the fluorescent structures. N indicates the number of structures analyzed for circularity. Data distributions are shown as combined violin and box plots. Statistical significance was calculated by Kruskal-Wallis test with post hoc Dunnett’s multiple comparisons, n.s.: nonsignificant, ***: *p* < 0.001, ****: *p* < 0.0001. (D) Flowchart of the *in vitro* reconstitution assay for TAS3-derived tasiRNA biogenesis used in this study. First, mRNAs encoding AGO7, SDE5, and *A. thaliana* SGS3 variants were added to the BY-2 lysate. Next, a synthetic miR390 duplex was introduced to assemble the AGO7–RISC complex. Endogenous *N. tabacum* SGS3 present in the BY-2 extract was then removed by immunodepletion. Finally, *TAS3* transcripts was added to trigger tasiRNA biogenesis. (E) *In vitro* tasiRNA biogenesis supported by SGS3 N-terminal region mutants. *TAS3* transcripts and 5′ D7(+) tasiRNA were detected by northern blotting, with U6 spliceosomal RNA used as a loading control. Representative results from two of six independent experiments are shown.

To dissect the functional contribution of the NCR, we generated three additional mutants:

1. Ala–EGFP, in which Asp/Glu residues between positions 161–190 were replaced with Ala;
2. Δ191–200–EGFP, lacking the region immediately C-terminal to the NCR, corresponding to a linker between the NCR and the zinc-finger domain; and (3) Δ161–200–EGFP, lacking both the entire NCR and the adjacent C-terminal linker region up to the zinc-finger domain (Figure 3A). Fluorescence microscopy revealed that the loss of the NCR resulted in a strong tendency to form irregular aggregates. Δ191–200–EGFP exhibited both small, spherical bodies and distorted aggregates, suggesting that the 191–200 region also contributes to preventing abnormal aggregation (Figure 3B). Quantitative analysis of the roundness of fluorescent structures observed in the nine constructs analyzed revealed three major groups (Figure 3C). The first group, including free-EGFP, ΔPrLDN–EGFP, and Δ160–EGFP, exhibited few distinct structures, with fluorescence signals diffusely distributed throughout the cytoplasm. The second group, represented by SGS3–EGFP and ΔZF–EGFP, exhibited well-defined, nearly spherical structures. The third group, including Δ200–EGFP, Δ161–200–EGFP, and Ala–EGFP, displayed irregular, non-circular assemblies distinct from the droplet-like structures observed in the wild type. Δ191–200–EGFP showed intermediate characteristics between the second and third groups, exhibiting both spherical structures and irregular assemblies; however, its roundness was significantly lower than that of the wild type.

To test whether the aggregation phenotype correlates with the ability to support tasiRNA biogenesis, we next examined the *in vitro* activity of these mutants using the cell-free assay. Mutants that formed irregular aggregates in *N. benthamiana*—Ala, Δ191–200, and Δ161–200—showed markedly reduced tasiRNA production compared with wild-type SGS3 (Figure 3D, E). Although the Δ191–200 mutant exhibited a substantial loss of tasiRNA production activity compared with the wild type, it retained partial activity relative to the Δ161–200 and Ala mutants, and the extent of tasiRNA accumulation varied among replicates.

Together, these observations reveal a close correlation between aberrant SGS3 assembly and impaired secondary siRNA biogenesis, raising the possibility that abnormal aggregation of SGS3 may directly compromise its functional competence in endogenous secondary siRNA production.

## Discussion

In this study, we dissected the molecular roles of the N-terminal IDR of SGS3 in secondary siRNA biogenesis. Using a combination of *in vitro* reconstitution, *in vivo* complementation, and imaging analyses, we showed that the PrLD of SGS3 is essential for siRNA body formation but dispensable for secondary siRNA production. Instead, a conserved negatively charged region (NCR) adjacent to the zinc-finger domain prevents abnormal aggregation of SGS3 in cells and is required for efficient tasiRNA biogenesis. These findings indicate that the N-terminal IDR of SGS3, through the actions of two distinct functional domains, regulates the aggregation propensity of SGS3 to ensure robust secondary siRNA amplification, while also controlling its LLPS-dependent subcellular localization independently of basal secondary siRNA production process.

Previous studies proposed that phase separation of SGS3 is required for secondary siRNA production (Tan *et al*, 2023; Kim *et al*, 2021). Our results, however, suggest an alternative interpretation. The deletion constructs used in earlier studies removed not only the N-terminal PrLD but also part of the adjacent NCR identified here. Loss of the NCR could have compromised SGS3 folding and/or solubility, resulting in impaired function independent of LLPS. Thus, our findings refine and redefine the current molecular model of secondary siRNA biogenesis by distinguishing the contribution of LLPS from that of the negatively charged, aggregation-preventing region within the N-terminal IDR (Figure 4).

**Figure 4.**
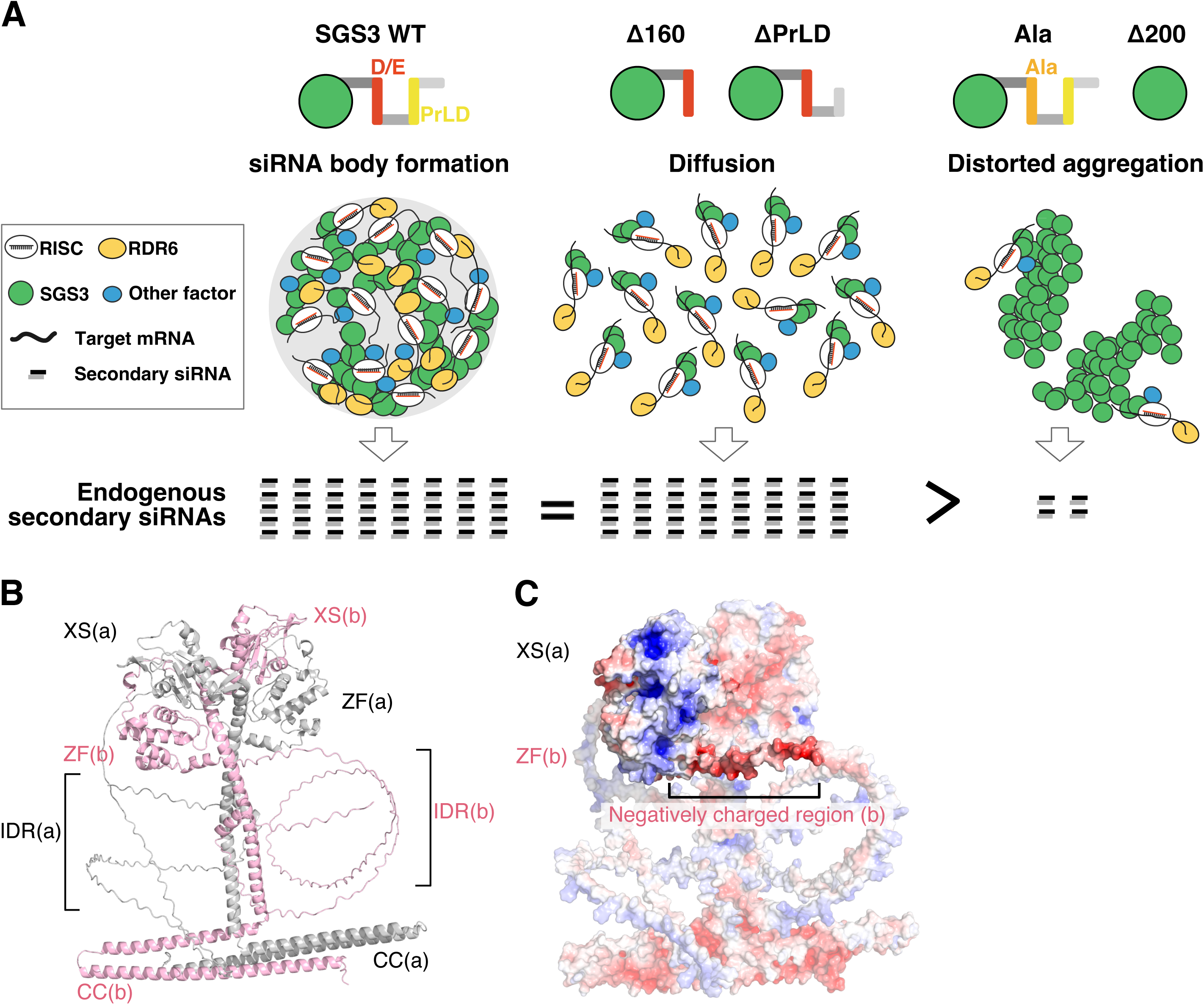
A functional model for the N-terminal region of SGS3. (A) Alanine substitution or deletion of the conserved NCR induces aberrant aggregation of SGS3 and results in defective tasiRNA production. In the presence of the NCR, the PrLD promotes LLPS and formation of siRNA bodies. However, siRNA body formation is not required for endogenous secondary siRNA biogenesis, which can be supported solely by the C-terminal 465 amino acids of SGS3. (B) Structural prediction of *Arabidopsis thaliana* SGS3 generated using AlphaFold 3. Consistent with previous reports (Fukunaga & Doudna, 2009), AtSGS3 forms a homodimer, with the two monomers shown in gray and pink. The negatively charged region (amino acids 161–190) is located adjacent to the ZF and XS domains. (C) A surface electrostatic potential map of the AtSGS3 structure. Blue indicates positively charged regions, whereas red represents negatively charged regions.

Structural prediction of SGS3, consistent with previous reports (Elmayan *et al*, 2009), indicates that SGS3 forms a homodimer and that the NCR is positioned adjacent to positively charged surface patches on the ZF domains and XS domains (Figure 4B, C). Because the XS domain is required for SGS3 to bind dsRNA (Fukunaga & Doudna, 2009), its positive charge is reasonable and likely reflects its functional role. The NCR may act to neutralize local positive charge of the ZF domains and XS domains, thereby preventing non-specific interactions with negatively charged nucleic acids or proteins. Indeed, negatively charged and intrinsically disordered regions have been reported in other systems to function as intrinsic anti-aggregation modules—for example, in histone chaperones, where they mimic DNA to neutralize positively charged surfaces and prevent non-native interactions (Warren & Shechter, 2017). Furthermore, consecutive Asp/Glu residues have also been reported to function as autonomous aggregation-suppressing tags in the context of recombinant protein purification (Jo, 2022; Su *et al*, 2007). Taken together with our observation that deletion of the negatively charged stretch leads to aberrant aggregation, it is reasonable to propose that, in SGS3, this NCR likewise prevents non-specific aggregation and maintains the protein in a functionally competent for specific recognition of RISC-targeted RNAs.

Our *in vivo* analyses further reveal that secondary siRNA biogenesis is remarkably robust to large variations in SGS3 abundance: tasiRNA accumulation changed only modestly even when SGS3 levels varied by approximately 100-fold across transgenic progeny (Figure 2). This argues strongly against a model in which LLPS-driven concentration of RISC, SDE5, or RDR6 is required for productive interactions. Instead, SGS3 likely acts through a stoichiometric, well-defined interaction with AGO–small RNA–target RNA complexes, where the number of accessible RISC–target sites—rather than total SGS3 concentration—limits RDR6 recruitment. Such a design allows the pathway to maintain stable output despite fluctuations in SGS3 abundance.

Although not required for basal tasiRNA production, the PrLD-mediated siRNA bodies may function as an inducible module that enhances silencing under high-demand conditions. During viral infection or bursts of transposon activity, local concentrations of target RNAs may increase dramatically; under such conditions, LLPS could accelerate RDR6 loading or dsRNA synthesis by spatially organizing SGS3-centered reaction hubs. Thus, LLPS may represent an adaptive amplification mechanism superimposed on a stoichiometric basal system.

In sum, our findings reveal that SGS3 uses two distinct IDR-embedded modules to coordinate activity and structural integrity: a PrLD that governs condensate formation, and a conserved NCR that prevents misfolding and aberrant aggregation. Together, these modules enable precise, robust, and condition-responsive control of secondary siRNA biogenesis. We propose that the N-terminal IDR acts as a molecular conductor—balancing flexibility with fidelity—to ensure that SGS3 engages productively with its partners and supports efficient amplification of small-RNA signals across diverse cellular contexts.

## Methods

### General methods

Lysate preparation from tobacco BY-2 cells and preparation of small RNA duplexes have been previously described in detail (Sakurai *et al*, 2021; Tomari & Iwakawa, 2017). *In vitro* tasiRNA biogenesis assay, immunodepletion of endogenous SGS3 from BY-2 lysate, and northern blot analyses used for the experiments shown in Figure 1 were performed as previously described (Sakurai *et al*, 2021). Recombinant AtSGS3 protein was expressed and purified as described previously (Iwakawa *et al*, 2021).

*In vitro* tasiRNA biogenesis assay and immunodepletion of endogenous SGS3 from BY-2 lysate conducted to obtain the data for Figure 3 involved slightly modified protocols, as described in below.

10 μL of BY-2 lysate, 4 μL of substrate mixture (250 µM each of 20 amino acids, 400 µM spermine, 125 mM creatine phosphate, 200 mM potassium acetate, 3.75 mM ATP, 0.5 mM GTP, 0.5 U/µL creatine phosphokinase (Sigma)), 2 μL of 1.5 μM miR390/ miR390* duplex, 1 μL of 1 μM 3× FLAG-*AtAGO7* mRNA, 0.5 µL of 1 μM 3×HA-*AtSDE5,* and 1 µL of 300 nM 3×HA-*AtSGS3* mRNA were mixed and incubated at 25 ℃. After120 min, 0.67 µg of anti-NtSGS3 was added and incubated on ice for 1h. To remove the antibodies and binding proteins thereof, the lysate was mixed with the pellet of 10 µL Dynabeads protein G (Invitrogen), and incubated at 4 °C for 1h. The supernatant was transferred into new tubes. Next, 0.4 μL of 100 nM *TAS3* RNAs was added to the supernatant and incubated at 25 °C for 30 min. Aliquots of the reaction mixtures were used for northern blotting after deproteinization with proteinase K (Nacalai Tesque, Inc.) and ethanol precipitation followed by dissolving the pellet in formamide dye (49% formamide, 5 mM ethylenediamine tetraacetic acid [EDTA], 0.01% xylene cyanol, and 0.01% bromophenol blue).

All primer and probe sequences used in this study are listed in Table S1.

### Northern Blot Assay

To detect U6 spliceosomal RNA and tasiRNAs, RNAs were separated on an 15% polyacrylamide denaturing gel and transferred onto a Nylon membranes, positively charged (Roche) using semidry transfer. The 1-ethyl-3- (3-dimethylaminopropyl) carbodiimide (EDC) cross-linking was used to fix RNA to the Nylon membrane, as described previously (57). The Nylon membrane was placed onto filter paper soaked in EDC solution and incubated at 60 °C for 1 h. To detect *TAS3* transcripts, a 6.7% polyacrylamide denaturing gel and a Hybond-N+ membrane (GE Healthcare) were used. RNAs were fixed to the Hybond-N+ membrane by ultraviolet cross-linking (120 mJ/cm^2^, 3 min). Membranes were placed in a hybridization bottle with 7.5 mL of PerfectHyb Plus Hybridization buffer (Sigma). The bottle was then preincubated for more than 5 min in the hybridization oven at 42 °C for U6, 68 °C for tasiRNA and *TAS3* transcripts, respectively. After preincubation, a probe, the sequences of the probes used and their corresponding detection targets are listed in Table S1, was hybridized with membranes overnight. The membrane was washed twice with low stringency buffer (2× saline-sodium citrate [SSC] and 0.1% SDS) at room temperature and then washed twice with high stringency buffer (0.1× SSC and 0.1% SDS) at the hybridization temperature for 15 min. Detection was performed according to the protocol for the DIG wash and block buffer set. Images were acquired using the Vilber Bio Imaging FUSION FX7. EDGE (M & S Instruments). DynaMarker, Prestain Marker for Small RNA Plus (BDL) was used as a marker.

### Preparation of Probes for Northern Blot Assay

For experiment involved in Figure 1, DNA oligo probes probe-1 and probe-2 were phosphorylated using T4 polynucleotide kinase (TaKaRa) with [γ- 32P]-ATP. probe-3, 4, 5, 6, 7, and 8 were mixed and radiolabeled as described for the detection of *TAS3* transcripts. For experiment involved in Figure 3, probe-9 and probe-10 were directly used. *TAS3* transcripts was detected with a Digoxigenin (DIG)-labeled long TAS3 probe, transcribed by using DIG RNA Labeling Kit (Roche). Template DNA was amplified from pUC57-TAS3a (Iwakawa *et al*, 2021) using oligo31 and oligo32 by using DIG RNA Labeling Kit (Roche).

### Quantification of SGS3 condensates in the cell-free system and Western blot assay

The mixture of tobacco BY-2 cell lysate and substrate mixture was supplemented to a final concentration of 45 nM with mRNA encoding EGFP, SGS3–EGFP, or ΔPrLD–EGFP. After incubation for 4 h at room temperature, 3 µL of each reaction was placed on a glass slide, covered with a coverslip, and imaged with a BZ-X810 microscope (KEYENCE) using a 10× objective, with the coverslip side facing downward. For each sample, two fields were acquired per reaction. Droplet numbers (“bodies”) were quantified in Fiji/ImageJ using ComDet v0.5.5 with the following parameters: Include larger particles = true, Segment larger particles = false, Approximate particle size = 5.0, and Intensity threshold (in SD) = 5.0. The mean of the two fields was taken as the value for each reaction. Three independent experiments were performed.

Simultaneously with microscopic observation, 2 µL of the sample was mixed with 18 µL of 1× SDS sample buffer (40% glycerol, 240 mM Tris·HCl pH 6.8, 8% SDS, 0.04% bromophenol blue, and dithiothreitol) and heat-denatured at 95 °C for 10 min. After electrophoresis, proteins were transferred onto a PVDF membrane (Merck) and incubated in blocking buffer (1% skim milk and 1× Tris buffered saline with Tween 20 [TBST]) for 30 min. Following blocking, the membrane was incubated overnight at room temperature with an anti-HA antibody (abcam, #ab130275) diluted 1:10,000 in blocking buffer. The membrane was then washed with TBST and incubated for 1 h at room temperature with anti-mouse immunoglobulin G (IgG)–horseradish peroxidase (HRP) conjugated antibodies (MBL, #330) diluted 1:25,000 in blocking buffer. After washing with TBST, the membrane was treated with a chemiluminescent substrate, and signals were detected and imaged using a FUSION FX7. EDGE (M & S Instruments)..

### Plasmid constructions

The following constructs used in this study have been previously described: pBYL2 (Mine *et al*, 2010), pBYL-3xFLAGAGO7 (Endo *et al*, 2013), pBYL-SGS3 (Sakurai *et al*, 2021), pBYL-3xFLAG-SGS3 (Sakurai *et al*, 2021), pEU-6xHis-SBP-SUMO-AtSGS3 (Iwakawa *et al*, 2021), pBYL-3xHA-SDE5 (Sakurai *et al*, 2021), pUC57-F-TAS3 (Iwakawa *et al*, 2021) and pUC57-TAS3a (Iwakawa *et al*, 2021). Primer sequences used in this study are listed in Table S1.

#### pEU-6xHis-SBP-SUMO-SGS3-3xHA

Two DNA fragments were prepared by PCR: SGS ORF fragment amplified from pEU-6xHis-SBP-SUMO-AtSGS3 (Iwakawa *et al*, 2021) using oligo21 and oligo28 and 3xHA tag sequence amplified using oligo29 and oligo30. The two DNA fragments were inserted into the SbfI/SalI-digested pEU-6xHis-SBP-SUMO-AtSGS3 vector (Iwakawa *et al*, 2021) using the HiFi DNA Assembly Cloning kit (New England Biolabs).

#### pEU-6xHis-SBP-SUMO-ΔN (40AA)-sgs3-3HA

Two DNA fragments were prepared by PCR: SGS ORF fragment amplified from pEU-6xHis-SBP-SUMO-AtSGS3-3xHA using primer pairs of oligo21-oligo35 and oligo36-oligo30. The two DNA fragments were inserted into the SbfI/SalI-digested pEU-6xHis-SBP-SUMO-AtSGS3-3xHA vector using the HiFi DNA Assembly Cloning kit (New England Biolabs).

#### pEU-6xHis-SBP-SUMO-ΔN (80AA)-sgs3-3HA

Two DNA fragments were prepared by PCR: SGS ORF fragment amplified from pEU-6xHis-SBP-SUMO-AtSGS3-3xHA using primer pairs of oligo21-oligo35 and oligo49-oligo30. The two DNA fragments were inserted into the SbfI/SalI-digested pEU-6xHis-SBP-SUMO-AtSGS3-3xHA vector using the HiFi DNA Assembly Cloning kit (New England Biolabs).

#### pEU-6xHis-SBP-SUMO-ΔN (120AA)-sgs3-3HA

Two DNA fragments were prepared by PCR: SGS ORF fragment amplified from pEU-6xHis-SBP-SUMO-AtSGS3-3xHA using primer pairs of oligo21-oligo35 and oligo50-oligo30. The two DNA fragments were inserted into the SbfI/SalI-digested pEU-6xHis-SBP-SUMO-AtSGS3-3xHA vector using the HiFi DNA Assembly Cloning kit (New England Biolabs).

#### pEU-6xHis-SBP-SUMO-ΔN (160AA)-sgs3-3HA

Two DNA fragments were prepared by PCR: SGS ORF fragment amplified from pEU-6xHis-SBP-SUMO-AtSGS3-3xHA using primer pairs of oligo21-oligo35 and oligo51-oligo30. The two DNA fragments were inserted into the SbfI/SalI-digested pEU-6xHis-SBP-SUMO-AtSGS3-3xHA vector using the HiFi DNA Assembly Cloning kit (New England Biolabs).

#### pEU-6xHis-SBP-SUMO-ΔN (200AA)-sgs3

The ORF of pEU-6xHis-SBP-SUMO-AtSGS3 (Iwakawa *et al*, 2021) was amplified by PCR using primer pairs oligo21 and oligo35, and oligo52 and oligo24. The resulting fragments were assembled into the pEU-6xHis-SBP-SUMO-AtSGS3 vector digested with SbfI and SalI using NEBuilder HiFi DNA Assembly Master Mix (NEB).

#### pEU-6xHis-SBP-SUMO-ΔN (200AA)-sgs3-3HA

The ORF of pEU-6xHis-SBP-SUMO-ΔN (200 AA)-sgs3 was amplified by PCR using primer pairs oligo21 and oligo28, and oligo29 and oligo30. The amplified fragments were assembled into the pEU-6xHis-SBP-SUMO-AtSGS3 vector digested with SbfI and SalI using NEBuilder HiFi DNA Assembly Master Mix (NEB).

#### pEU-6xHis-SBP-SUMO-ΔPrLD_N-sgs3-3HA

Three DNA fragments were prepared by PCR: SGS ORF fragment amplified from pEU-6xHis-SBP-SUMO-AtSGS3-3xHA using primer pairs of oligo21-oligo31, oligo32-oligo28 and oligo30-oligo31. The three DNA fragments were inserted into the SbfI/SalI-digested pEU-6xHis-SBP-SUMO-AtSGS3-3xHA vector using the HiFi DNA Assembly Cloning kit (New England Biolabs).

#### pEU-6xHis-SBP-SUMO-ΔPrLD_NC-sgs3-3HA

Four DNA fragments were prepared by PCR: SGS ORF fragment amplified from pEU-6xHis-SBP-SUMO-AtSGS3-3xHA using primer pairs of oligo21-oligo31, oligo32-oligo33, oligo34-oligo28 and oligo30-oligo31. The four DNA fragments were inserted into the SbfI/SalI-digested pEU-6xHis-SBP-SUMO-AtSGS3-3xHA vector using the HiFi DNA Assembly Cloning kit (New England Biolabs).

#### pEU-6xHis-SBP-SUMO-ΔPrLD_C-sgs3-3HA

Three DNA fragments were prepared by PCR: SGS ORF fragment amplified from pEU-6xHis-SBP-SUMO-AtSGS3-3xHA using primer pairs of oligo21-oligo33, oligo34-oligo28 and oligo30-oligo31. The four DNA fragments were inserted into the SbfI/SalI-digested pEU-6xHis-SBP-SUMO-AtSGS3-3xHA vector using the HiFi DNA Assembly Cloning kit (New England Biolabs).

#### pEU-6xHis-SBP-SUMO-sgs3-E500K

The 1498th base in the ORF of SGS3 was changed from Guanine to Adenine from pEU-6xHis-SBP-SUMO-AtSGS3 (Iwakawa *et al*, 2021) by site-directed mutagenesis PCR by using oligo47 and oligo48.

#### pEU-6xHis-SBP-SUMO-sgs3-ΔZF

Two DNA fragments were prepared by PCR: SGS ORF fragment amplified from pEU-6xHis-SBP-SUMO-AtSGS3 (Iwakawa *et al*, 2021) using primer pairs oligo21-oligo22 and oligo23-oligo24. The two DNA fragments were inserted into the SbfI/SalI-digested pEU-6xHis-SBP-SUMO-AtSGS3 vector (Iwakawa *et al*, 2021) using the HiFi DNA Assembly Cloning kit (New England Biolabs).

#### pEU-6xHis-SBP-SUMO-sgs3-ΔXS

Two DNA fragments were prepared by PCR: SGS ORF fragment amplified from pEU-6xHis-SBP-SUMO-AtSGS3 (Iwakawa *et al*, 2021) using primer pairs oligo21-oligo25 and oligo23-oligo26. The two DNA fragments were inserted into the SbfI/SalI-digested pEU-6xHis-SBP-SUMO-AtSGS3 vector (Iwakawa *et al*, 2021) using the HiFi DNA Assembly Cloning kit (New England Biolabs).

#### pEU-6xHis-SBP-SUMO-sgs3-ΔCC

The SGS ORF fragment was amplified from pEU-6xHis-SBP-SUMO-AtSGS3 (Iwakawa *et al*, 2021) using primer pairs oligo21-oligo27. This DNA fragments was inserted into the SbfI/SalI-digested pEU-6xHis-SBP-SUMO-AtSGS3 vector (Iwakawa *et al*, 2021) using the HiFi DNA Assembly Cloning kit (New England Biolabs).

#### pBYL_3xHA_EGFP

Two DNA fragments were prepared by PCR: 3xHA tag sequence amplified using oligo13 and oligo14, EGFP sequence amplified using oligo15 and oligo16. The two DNA fragments were cloned into the XbaI/AscI-digested pBYL-SGS3 vector (Sakurai *et al*, 2021) using the HiFi DNA Assembly Cloning kit (New England Biolabs).

#### pBYL_3xHA_SGS3_EGFP

Three DNA fragments were prepared by PCR: 3xHA tag sequence amplified using oligo17 and oligo19, SGS ORF fragment amplified from pBYL-SGS3 (Sakurai *et al*, 2021) using oligo18 and oligo20 and EGFP sequence amplified using oligo15 and oligo16. The three DNA fragments were inserted into the MluI/AscI-digested pBUL_3xHA_EGFP vector via HiFi DNA Assembly Cloning kit (New England Biolabs).

#### pBYL_3xHA_ΔPrLD_N_sgs3_EGFP

Three DNA fragments were prepared by PCR: 3xHA tag sequence amplified using oligo17 and oligo19, SGS ORF fragment amplified from *pEU-6xHis-SBP-SUMO-ΔPrLD_N -sgs3-3HA* using oligo18 and oligo20 and EGFP sequence amplified using oligo15 and oligo16. The three DNA fragments were inserted into the MluI/AscI-digested pBUL_3xHA_EGFP vector via HiFi DNA Assembly Cloning kit (New England Biolabs).

#### pEarleyGate_3xFLAG_SGS3

Two DNA fragments were prepared by PCR: 3xFLAG tag sequence amplified using oligo38 and oligo37 from pBYL-3xFLAGAGO7, SGS ORF fragment amplified from pBYL-3xFLAG-SGS3 (Sakurai *et al*, 2021) using oligo39 and oligo40. The two DNA fragments were cloned into the XhoI/XbaI-digested pEarleyGate 100 vector (Earley *et al*, 2006) using the HiFi DNA Assembly Cloning kit (New England Biolabs).

#### pEarleyGate_3xFLAG_ΔPrLD_N_sgs3

Two DNA fragments were prepared by PCR: 3xFLAG tag sequence amplified using oligo38 and oligo37 from pBYL-3xFLAGAGO7, ΔPrLD_N_sgs3 ORF fragment amplified from pEU-6xHis-SBP-SUMO-ΔPrLD_N-sgs3-3HA using oligo39 and oligo40. The two DNA fragments were cloned into the XhoI/XbaI-digested pEarleyGate 100 vector using the HiFi DNA Assembly Cloning kit (New England Biolabs).

#### pEarleyGate_3xFLAG_ΔN (160AA)-sgs3

Two DNA fragments were prepared by PCR: 3xFLAG tag sequence amplified using oligo38 and oligo37 from pBYL-3xFLAGAGO7, ΔN (160AA)-sgs3 ORF fragment amplified from pEU-6xHis-SBP-SUMO-ΔN (160AA)-sgs3-3HA using oligo41 and oligo40. The two DNA fragments were cloned into the XhoI/XbaI-digested pEarleyGate 100 vector using the HiFi DNA Assembly Cloning kit (New England Biolabs).

#### pEarleyGate_3xFLAG_ΔN (200AA)-sgs3

Two DNA fragments were prepared by PCR: 3xFLAG tag sequence amplified using oligo38 and oligo37 from pBYL-3xFLAGAGO7, ΔN (160AA)-sgs3 ORF fragment amplified from pEU-6xHis-SBP-SUMO-ΔN (200AA)-sgs3-3HA using oligo42 and oligo40. The two DNA fragments were cloned into the XhoI/XbaI-digested pEarleyGate 100 vector using the HiFi DNA Assembly Cloning kit (New England Biolabs).

#### pEarleyGate_3xFLAG_Fluc

Two DNA fragments were prepared by PCR: 3xFLAG tag sequence amplified using oligo38 and oligo37 from pBYL-3xFLAGAGO7, Firefly luciferase fragment amplified from psiCHECK^TM^-2 vector (Promega) using oligo43 and oligo44. The two DNA fragments were cloned into the XhoI/XbaI-digested pEarleyGate 100 vector using the HiFi DNA Assembly Cloning kit (New England Biolabs).

#### pENTA-3xHA _EGFP

In order to obtain the vector sequence, DNA fragments were amplified from the entry vector pENTA (Himeno *et al*, 2010) using oligo1 and oligo2. 3xHA _EGFP ORF fragment amplified from pBYL-3xHA _EGFP using oligo3 and oligo4. The two DNA fragments were assembled via HiFi DNA Assembly Cloning kit (New England Biolabs).

#### pENTA-3xHA_SGS3_EGFP

In order to obtain the vector sequence, DNA fragments were amplified from the entry vector pENTA using oligo1 and oligo2. 3xHA_SGS3_EGFP ORF fragment amplified from pBYL-3xHA_SGS3_EGFP using oligo3 and oligo4. The two DNA fragments were assembled via HiFi DNA Assembly Cloning kit (New England Biolabs).

#### pENTA-3xHA_ΔPrLD_SGS3_EGFP

In order to obtain the vector sequence, DNA fragments were amplified from the entry vector pENTA using oligo1 and oligo2. 3xHA_ΔPrLD_SGS3_EGFP ORF fragment amplified from pBYL-3xHA_ΔPrLD_SGS3_EGFP using oligoF3 and oligoF4. The two DNA fragments were assembled via HiFi DNA Assembly Cloning kit (New England Biolabs).

#### pENTA-3xHA_Δ160_SGS3_EGFP

In order to obtain the vector sequence, DNA fragments were amplified from the entry vector pENTA using oligo1 and oligo2. 3xHA_Δ160_SGS3_EGFP ORF fragment amplified from pBYL-3xHA_Δ160_SGS3_EGFP using oligo3 and oligo4. The two DNA fragments were assembled via HiFi DNA Assembly Cloning kit (New England Biolabs).

#### pENTA-3xHA_Δ200_SGS3_EGFP

In order to obtain the vector sequence, DNA fragments were amplified from the entry vector pENTA using oligo1 and oligo2. 3xHA_Δ200_SGS3_EGFP ORF fragment amplified from pBYL-3xHA_Δ200_SGS3_EGFP using oligo3 and oligo4. The two DNA fragments were assembled via HiFi DNA Assembly Cloning kit (New England Biolabs).

#### pENTA-3xHA_Δ161-200_SGS3_EGFP

DNA fragments were amplified from pENTA-3xHA_SGS3_EGFP using oligo5 and oligo6. DNA fragment was directly introduced to *E. coli* competent cells.

#### pENTA-3xHA_ΔNCP_SGS3_EGFP

DNA fragments were amplified from pENTA-3xHA_SGS3_EGFP using oligo9 and oligo10. DNA fragment was directly introduced to *E. coli* competent cells.

#### pENTA-3xHA_Δ191-200_SGS3_EGFP

DNA fragments were amplified from pENTA-3xHA_SGS3_EGFP using oligo7 and oligo8. DNA fragment was directly introduced to *E. coli* competent cells.

*pEarleyGate-3xHA_EGFP, pEarleyGate-3xHA_SGS3_EGFP, pEarleyGate-3xHA_ΔPrLD_SGS3_EGFPΔPrLD, pEarleyGate-3xHA_Δ160_SGS3_EGFP, pEarleyGate-3xHA_Δ200_SGS3_EGFP, pEarleyGate-3xHA_Δ161-200_SGS3_EGFP, pEarleyGate-3xHA_ΔNCP_SGS3_EGFP, pEarleyGate-3xHA_Δ191-200_SGS3_EGFP*

pENTA-3xHA_EGFP, pENTA-3xHA_SGS3_EGFP, pENTA-3xHA_ΔPrLD_SGS3_EGFP, pENTA-3xHA_Δ160_SGS3_EGFP, pENTA-3xHA_Δ200_SGS3_EGFP, pENTA-3xHA_Δ 161-200_SGS3_EGFP, pENTA-3xHA_ΔNCP_SGS3_EGFP, and pENTA-3xHA_Δ191-200_SGS3_EGFP plasmids were inserted into pEarleyGate 100 vector by using LR Clonase (Invitrogen) reaction-mediated recombination system.

### *In vitro* transcription

mRNAs were transcribed *in vitro* from NotI- (for plasmids with the prefix ‘‘pBYL-’’) or XhoI-(for plasmids with the prefix ‘‘pUC57-’’) digested plasmids or PCR products amplified from pENTA-plasmids by using Oligo11 and Oligo12 primer set using the AmpliScribe T7 High Yield Transcription Kit (Lucigen) or T7-Scribe^TM^ Standard RNA IVT Kit (Cell Script), followed by capping with ScriptCap m7G Capping System (Cell Script). Poly(A)-tails were added to the transcripts by the A-Plus Poly(A) Polymerase Tailing Kit (Cell Script).

### Plants transformation and transient expression

*A. thaliana* Col-0 plants were transformed with *Agrobacterium tumefaciens* EHA105 harboring pEarleyGate-3xFLAG_Fluc or its variants, pEarleyGate-3xFLAG_SGS3, pEarleyGate-3xFLAG_ΔPrLDN_SGS3, pEarleyGate-3xFLAG_Δ160_SGS3, and pEarleyGate-3xFLAG_Δ200_SGS3. T1 transgenic seeds were sown on soil mixed with Supermix (Sakata) and vermiculite (Nittai) (1:1) and placed at 4 °C for 3–5 d before transfer to growth chambers under 16-h day/8-h night illumination cycles with a temperature of 22 °C. Flower buds of BASTA-resistant T1 plants were harvested in liquid nitrogen and stored at -80°C. Frozen flower buds were ground to powder using a mortar and pestle and approximately one spatula scoop of the powder was mixed with 100 µL of 1.5 x SDS-PAGE sample buffer and heat-denatured at 95 °C for 5 min. To detect FLAG-tagged SGS3 anti-DYKDDDDK tag (WAKO 012-22384), goat anti-mouse IgG (H+L) (Jackson ImmunoResearch 115-035-003) and Pierce ECL Plus Western Blotting Substrate (Thermo Fisher Scientific 32132). RNA was extracted with TRIzol Reagent (Invitrogen 15596-018) from remaining powder that was further disrupted using a bead beater and glass beads.

*Nicotiana* plants were grown under long day (16 hours light:8 hours dark) conditions at 25 ℃. Agroinfiltration was performed as previously described (Llave *et al*, 2000). The Agrobacterium tumefaciens GV3101 (pMP90) (Koncz & Schell, 1986) transformed with pEarleyGate-3xHA_EGFP or its variants, pEarleyGate-3xHA_SGS3_EGFP, pEarleyGate-3xHA_Δ PrLD_SGS3_EGFP, pEarleyGate-3xHA_Δ160_SGS3_EGFP, pEarleyGate-3xHA_Δ 200_SGS3_EGFP, pEarleyGate-3xHA_Δ161-200_SGS3_EGFP, pEarleyGate -3xHA_Δ NCP_SGS3_EGFP, and pEarleyGate-3xHA_Δ191-200_SGS3_EGFP were pooled (optical density at 600 nm (OD600) = 0.2). The Agrobacterium suspension was infiltrated into leaves using a 1 mL syringe without needle. The leaves were harvested at 48 h post-infiltration. *Nicotiana* leaves were directly imaged on a Zeiss LSM980 (Zeiss).

All images were analyzed using a semi-automated Fiji-based macro designed to quantify phase-separated droplets in *Nicotiana* leaves. The macro measures “Area”, “Perimeter”, “Circularity”, “Major Axis Length”, and “Minor Axis Length”. Roundness was calculated using the following formula: Roundness = 4×Area/ π×(Major Axis Length)^2

The statistical analyses were performed using R statistical software, version 4.4.2. Dunnett’s test was used for multiple comparisons, and p-values were adjusted for multiple comparisons using the Holm method.

### Multiple sequence alignment

The SGS3 sequences of 11 different kinds of plant (*Arabidopsis thaliana*, *Glycine max*, *Solanum lycopersicum*, *Cucurbita moschata*, *Erythranthe lewisii*, *Nelumbo nucifera*, *Rhizophora mucronata*, *Coffea arabica*, *Eucalyptus grandis*, *Actinidia chinensis* var. chinensis, *Hibiscus syriacus* and *Marchantia polymorpha*) were aligned using the Jalview of Clustal Omega (Sievers *et al*, 2011). The sequence of each kinds of SGS3 were obtained from the database from Uniprot website (https://www.uniprot.org/docs/pkinfam) or NCBI website (https://www.ncbi.nlm.nih.gov).

### Structure prediction and visualization of AtSGS3

The structure prediction of the SGS3 homodimer was performed using the AlphaFold 3 (Abramson *et al*, 2024) (https://alphafoldserver.com/) . Structural visualization and comparative analysis were carried out using PyMOL (Schrödinger).

### Small RNA sequence Analysis

Adapters were trimmed as supplied by the provider. Reference data were prepared from *A. thaliana* TAIR10 (whole genome and transcript models) (Berardini *et al*, 2015). Transcript fasta sequences were extracted with gffread (Pertea & Pertea, 2020). Mature *A. thaliana* miRNAs were obtained from miRBase (Kozomara *et al*, 2019).

To remove miRNA reads, raw small-RNA reads were first aligned to miRBase with Bowtie (-a - -best --strata -v 2, ≤2 mismatches) (Langmead *et al*, 2009). Mapped read IDs were excluded (“miRNA-depletion”), and the remaining reads were used for all downstream analyses. miRNA-depleted reads were then aligned with perfect matching (-v 0) to the TAIR10 genome and transcript references. SAM/BAM files were converted to BED format with sam tools and BEDTools (Li *et al*, 2009; Quinlan & Hall, 2010). For each library and reference, coverage was computed using bedtools coverage.

Strand-specific 21-nt coverage profiles were generated for TAS1–3 loci by mapping miRNA-depleted reads to TAIR10 transcripts with perfect matching and extracting reads corresponding to TAS gene names. Coverage values were normalized to total genome-mapped small-RNA reads (RPM) before plotting strand-specific distributions. Graphs were generated from these normalized data in R using Biostrings (Pagès *et al*, 2025), ggplot2 (Wickham, 2016) and base graphics.

## Data availability

The sequencing data reported in this paper are publicly available in DDBJ, under the accession number DRR801829-DRR801840.

## Acknowledgements

We are also grateful to all the members of the Iwakawa laboratory and Tomari laboratory for fruitful discussion; and Yukihide Tomari for critically reading the manuscript. This work was supported by JSPS KAKENHI (grant numbers 23H02412 to H.-o.I., 22KJ2862 to Y.F. and 20J11529 to Y.S.), JST FOREST (grant JPMJFR204O to H.-o.I.), JST PRESTO (grant JPMJPR18K2 to H.-o.I.).

## Author Contributions

Y.F., Y.S., and H.-o.I. designed the project. Y.F., Y.S., M.Y., and H.-o.I. performed biochemical experiments. K.S. performed bioinformatic analyses. H.-o.I. supervised the project. Y.F., Y.S., R.K., K.S., M.Y., and H.-o.I. analyzed the data and wrote the manuscript. All authors discussed the results and approved the final version of the manuscript.

## Conflict of Interest

The authors declare no competing interests.

## Extended view figure legends

**Figure EV. 1.**
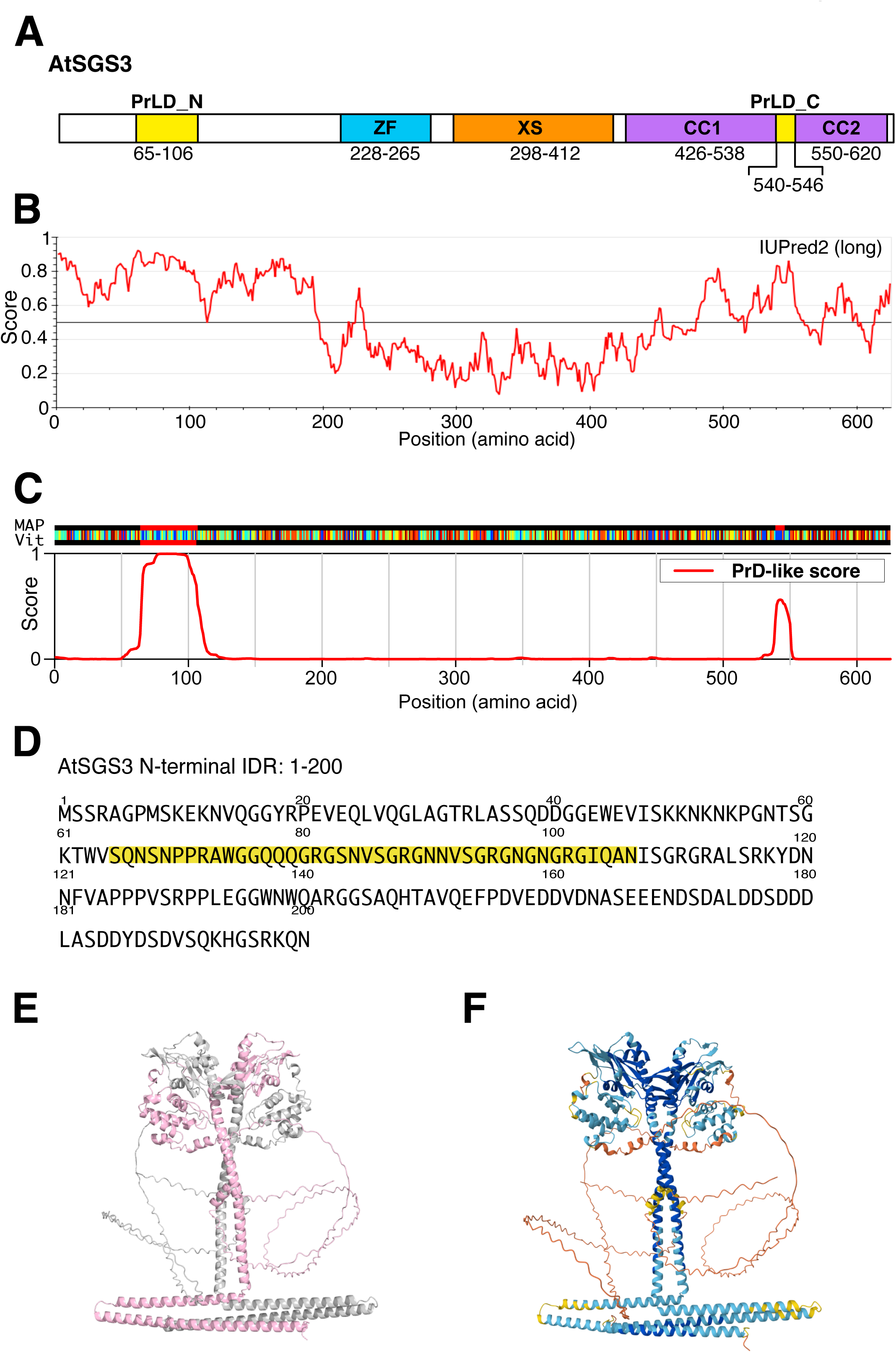
SGS3 possesses an intrinsically disordered region (IDR) at its N terminus that contains a prion-like domain. (A) Predicted domain organization of *A. thaliana* SGS3. SGS3 is predicted to contain a ZF domain, an XS domain, and a coiled-coil (CC) domain. (B) Prediction of intrinsically disordered regions (IDRs) in SGS3 using the IUPred2 algorithm (Mészáros *et al*, 2018; Erdős & Dosztányi, 2020). Residues with IUPred scores above 0.5 are likely to be part of intrinsically disordered regions. Notably, the ZF and XS domains correspond to regions with IUPred scores below 0.5, suggesting that they form stable, structured domains. (C) Prediction of PrLDs in SGS3 using the PLAAC algorithm (Lancaster *et al*, 2014). Regions with scores close to 1 indicate amino acid compositions similar to those of PrLDs. Two PrLDs were predicted within SGS3: one spanning residues 65–106 and another from residues 540–546. (D) Amino acid sequence of the N-terminal 200 residues of SGS3. The PrLD region is highlighted in yellow. (E, F) Structural prediction of *A. thaliana* SGS3 generated using AlphaFold 3. SGS3 is predicted to formed a homodimer. In the figure, the two molecules are shown in different colors—gray and pink—for clarity in (E). A per-residue confidence score map is shown in (F), with colors indicating the confidence level: dark blue represents the highest confidence, followed by light blue, yellow, and orange as the confidence decreases. The N-terminal region is predicted to be structurally flexible and difficult to model.

**Figure EV. 2.**
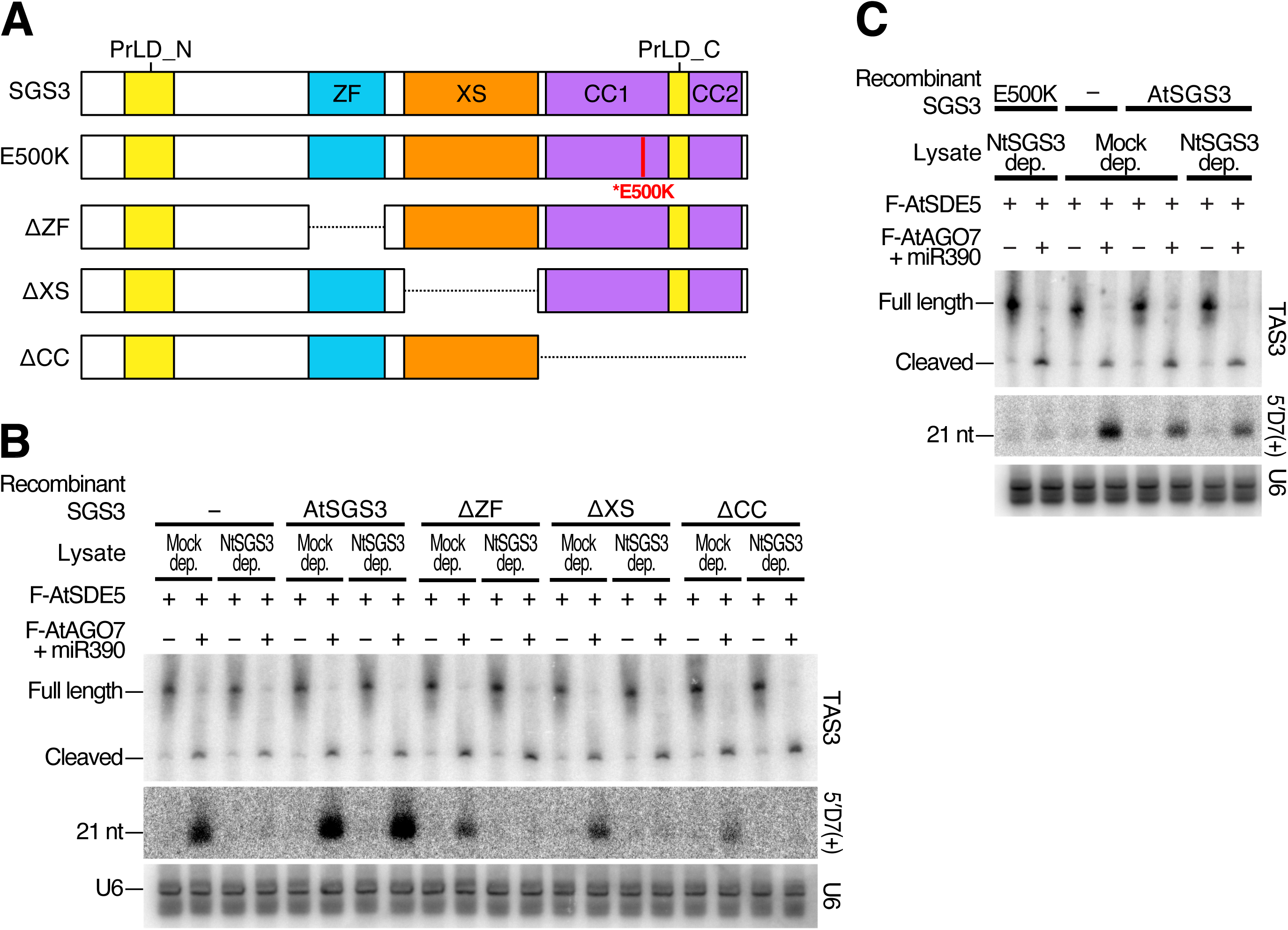
The previously identified functional domains (ZF, XS, and CC domains) and key amino acid residues of SGS3 are essential for tasiRNA biogenesis *in vitro*. (A) Schematic representation of SGS3 mutants with alterations in known functional domains or critical amino acid residues. The second construct from the top represents the sgs3-E500K mutant, in which a point mutation at the 500th amino acid residue of SGS3 negatively affects tasiRNA production, as previously reported in studies using *Arabidopsis* plants (Adenot *et al*, 2006). The red mark indicates the position of this mutation, which is located within the CC1 domain. The bottom three constructs represent deletion mutants lacking the predicted domain regions (sgs3-ΔZF, sgs3-ΔXS, and sgs3-ΔCC). Dashed lines indicate deleted regions. (B, C) In vitro tasiRNA biogenesis with SGS3 mutants shown in (A). *TAS3* transcripts and 5′ D7(+) tasiRNA were detected by northern blotting, with U6 spliceosomal RNA serving as a loading control. Representative results from two of six independent experiments are shown.

**Figure EV. 3.**
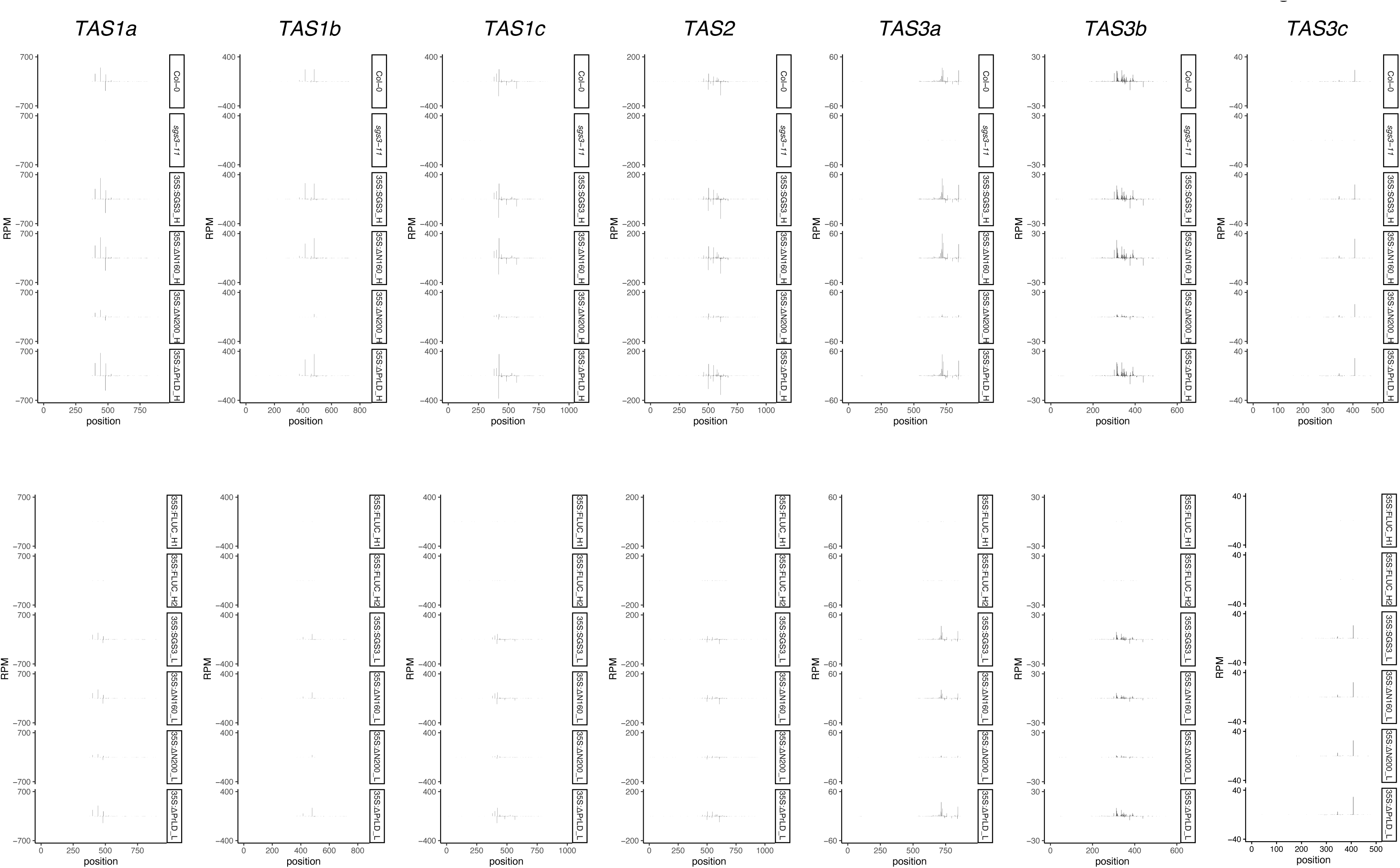
5′-end mapping profiles of 21-nt tasiRNAs at *TAS1*, *TAS2*, and *TAS3* loci in Col-0, sgs3-11, and SGS3-complemented lines. Profiles of 21-nt tasiRNAs mapped along the genomic regions of *TAS1a, TAS1b*, *TAS1c*, *TAS2*, *TAS3a*, *TAS3b*, and *TAS3c*, plotted according to the 5′ ends of small RNA reads (RPM). For each locus, profiles from wild-type Col-0, sgs3-11, and sgs3-11 progeny complemented with WT SGS3 or SGS3 mutant constructs (high-expression “H” and low-expression “L”) are shown. The x-axis represents the genomic coordinates of each TAS locus (including introns where present), and the y-axis indicates the abundance of 5′-end small RNA reads in reads per million (RPM).

**Figure EV. 4.**
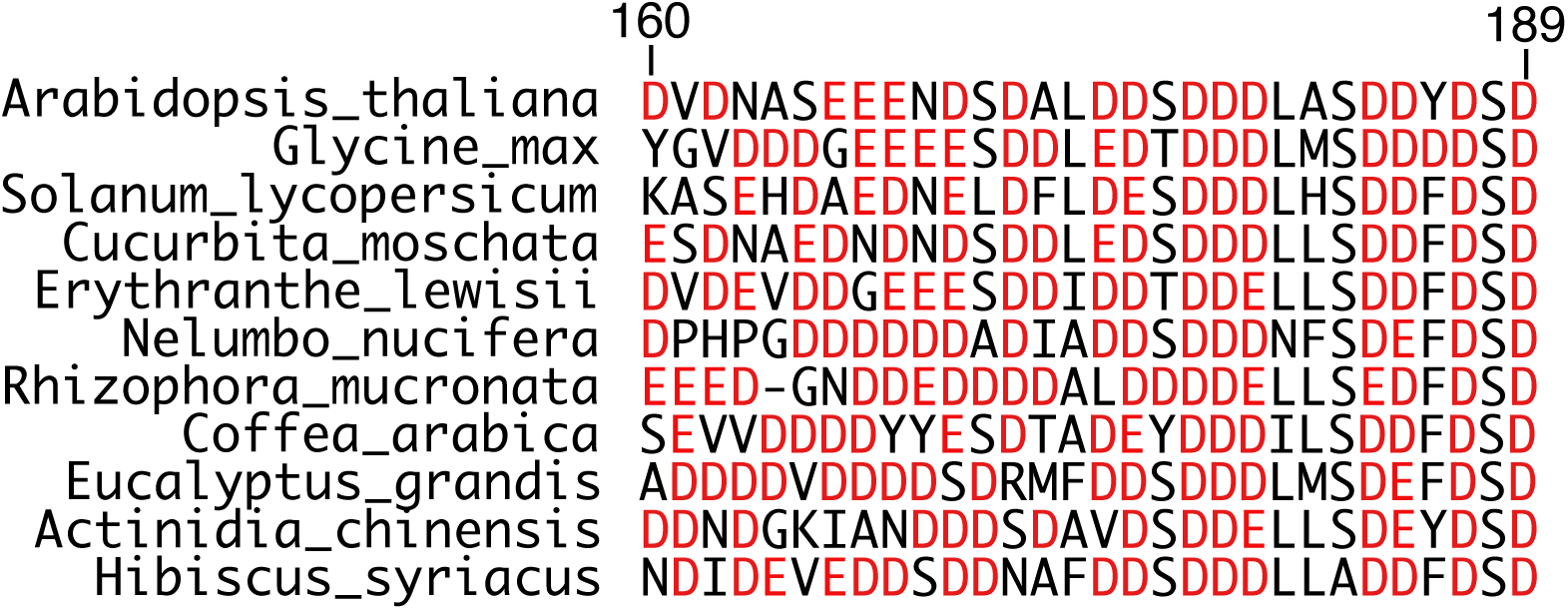
Conservation of the N-terminal negatively charged region of SGS3 among different angiosperm species. Amino acid sequences of the 160–189 region of *Arabidopsis* SGS3 and the corresponding regions of SGS3 from other plant species are shown. Numbers indicate amino acid positions in *Arabidopsis* SGS3. The alignment was performed using Clustal Omega (Sievers *et al*, 2011).

